# Spatial localisation of touch on a robotic limb in the absence of direct haptic feedback

**DOI:** 10.64898/2026.06.29.735244

**Authors:** Celia Foster, Martina Giancane, Valeria C. Peviani, Andrew Dott, Eva Chapman, Mario Kleiner, Luke E. Miller, Calogero Maria Oddo, Dani Clode, Tamar R. Makin

## Abstract

Artificial limbs typically lack somatosensory receptors, limiting users’ access to haptic feedback about movement outcomes. However, indirect tactile signals generated during object interaction are transmitted to the body-limb interface and may serve as a haptic feedback source. This study investigates whether such indirect tactile signals convey distinct information that can support localisation of touch on an artificial body part. Using an additional robotic thumb (The Third Thumb, Dani Clode Design), we measured tactile signals at the interface between the Third Thumb and the hand following stimulation from four vibration motors positioned along the Thumb. We then assessed whether participants could discriminate these signals and integrate them with different Thumb postures to spatially localise touch. Measurements showed that each stimulation site produced distinct tactile signatures at the interface. Behaviourally, participants localised touch at above-chance levels. Representational similarity analyses further revealed that performance was best explained by the spatial positions of stimuli, rather than by limb-based coordinates or indirect signal patterns alone, indicating flexible mapping of indirect touch signals onto the Third Thumb posture. Together, these findings demonstrate that indirect tactile signals transmitted through an artificial limb can be discriminated and flexibly remapped to support spatial localisation of touch.

## 1. Introduction

Sensory feedback following movement is widely recognised as fundamental to motor control and learning (Wolpert et al., 1995). In particular, haptic and proprioceptive inputs play a crucial role, as evidenced by profound motor impairments observed in deafferented patients who lack tactile and proprioceptive afferent signals despite retaining intact motor output pathways (Cole, 1995; Richardson et al., 2016). Deficits in motor ability have also been reported under experimental conditions in which somatosensory input is transiently disrupted, such as through the use of anaesthetic nerve blocks (Carteron et al., 2016). Haptic feedback provided during the sensorimotor control loop has been shown to underlie perception of object texture and materiality and support stable grasping (Klatzky, 2025). Together, these findings highlight the essential contribution of haptic feedback to effective motor control.

Robotic limbs are designed both to restore motor function in clinical populations and to extend movement capabilities through augmentation. Achieving this requires effective integration with the motor system to enable movement, as well as with the somatosensory system to deliver haptic feedback about movement outcomes (Dowdall, Foster, et al., 2025; Nisky & Makin, 2024). In the context of restorative artificial limbs, interfaces with the somatosensory system have been developed at multiple levels, including peripheral and central nerves, the skin, and the brain, and these approaches have been shown to improve motor control in amputee users (Bensmaia et al., 2020; Roche et al., 2023; Shokur et al., 2026). In contrast, for augmentative robotic limbs that aim to introduce entirely new effectors, it remains unclear how best to integrate artificial sensory signals with the existing somatosensory system (Dowdall, Foster, et al., 2025). One commonly explored approach is somatosensory substitution, in which sensory signals from the artificial limb are remapped to an alternative skin site (Hussain & Prattichizzo, 2020). Although this method can enhance control of additional effectors, it is constrained by the repurposing of skin areas already dedicated to natural sensation and by the ongoing cognitive effort required to reinterpret remapped inputs. Consequently, current non-invasive, skin-based methods for delivering haptic feedback present significant limitations, particularly for augmentative applications (Dowdall, Foster, et al., 2025).

An emerging approach for delivering haptic feedback in wearable robotic limbs is to exploit indirect touch signals that are naturally transmitted to the interface between the robotic limb and the user’s body. Indirect touch signals arise when the robotic limb moves or makes contact with objects and are conveyed through the structure of the limb interface of the wearable robotic limb with the users’ body. Initial work demonstrated that indirect touch signals can enhance motor control of a wearable robotic limb interfacing with the torso of the user (Guggenheim & Asada, 2021; Parietti & Asada, 2017). Subsequent studies have shown that indirect touch signals can convey rich sensory information, enabling users to discriminate object and texture properties perceived via an augmentative robotic limb (Dowdall, Molina-Sanchez, et al., 2025). Indirect touch has also been shown to indicate impact location on a prosthetic limb (Ivani et al., 2024), and early pioneering work demonstrated that coupling of distal muscles to the movement of a prosthetic limb could provide proprioceptive feedback (Doubler & Childress, 1984). Following training with a augmentative robotic limb, participants exhibit improved non-visual motor coordination, suggesting that these signals can support inference of the limb’s posture (Kieliba et al., 2021; Molina-Sanchez et al., 2026). Complementary research on handheld tools has indicated that humans can localise the point of impact on a tool with high accuracy using the same principle of indirect touch (Miller et al., 2018; Peviani et al., 2026). However, such studies have tested this ability with only rigid tools made out of a single consistent material. It remains unclear whether such localisation abilities generalise to robotic limbs, with curved or more complex shape that changes with posture and heterogeneous material properties. Establishing this would be critical for determining whether indirect touch can provide precise information about the location of object interactions on wearable robotic limbs.

Localising touch spatially relies on the integration of tactile signals with information about current body posture. Tactile inputs are initially encoded in cortex in a skin-based reference frame, reflecting the spatial arrangement of mechanoreceptors across the body surface (Penfield & Boldrey, 1937; Roux et al., 2018). However, because body posture is dynamic, this skin-based information must be transformed into a spatial reference frame to determine the location of a tactile stimulus in space (Klautke et al., 2023), a process termed tactile remapping. Body posture information is determined using multisensory integration of proprioception and vision (van Beers et al., 1999). Consequently, visual information alone can modulate tactile perception and remapping, for example, using visual input from a rubber hand, when the real hands are occluded (Azañón & Soto-Faraco, 2007; Botvinick & Cohen, 1998; Shore & Cadieux, 2013). Furthermore, congenitally blind individuals exhibit marked differences in tactile remapping compared to both sighted and late-blind individuals (Röder et al., 2004, 2007).

Temporal-order judgement paradigms reveal systematic performance deficits when skin-based and external spatial reference frames are misaligned, for example, when the limbs are crossed (Heed & Azañón, 2014; Yamamoto & Kitazawa, 2001a). Notably, similar effects have been observed when rigid handheld tools are crossed while the biological limbs remain uncrossed (Yamamoto et al., 2005; Yamamoto & Kitazawa, 2001b). This and other related findings (Miller et al., 2023; Themelis et al., 2026) suggest that indirect touch signals can be remapped to the external spatial position of tools for hand extension (e.g. sticks). However, the limits of tactile remapping remain unclear, in particular, whether touch can be remapped to artificial limbs with complex, posture-dependant morphology and augmentative artificial limbs that add to the body - fundamentally altering body morphology. Furthermore, it is unclear whether tactile remapping depends on motor experience, and consequently whether such experience is required for successful remapping. Addressing these questions is critical for determining whether indirect touch can support functional haptic feedback in augmentative artificial limb systems.

The present study aimed to determine whether indirect touch signals provide distinct information indicating touch location on a robotic limb, and whether human participants can discriminate these signals and remap them to the limb’s current posture. To address these questions, we used the Third Thumb (Dani Clode Design) a wearable, additional robotic thumb with two degrees of freedom (**Figure 1A**). The Third Thumb attaches to the hand via a handpiece positioned on the lateral side of the palm below the little finger. For the Third Thumb, indirect touch signals arise from contact with external objects or movements of the Third Thumb, which are transmitted through the robotic limb to the skin beneath the handpiece. For the present study, we developed a modified version of the Third Thumb with vibration motors placed at four locations to deliver touch stimulation at these locations (**Figure 1B**). In addition, we selected six distinct Third Thumb postures, corresponding to six equal steps along the first degree of freedom (flexion – extension) of the Third Thumb, without any contact of the Third Thumb tip with the palm. Together, these manipulations allowed us to systematically investigate both the discrimination of indirect touch signals and their transformation relative to the limb’s spatial posture.

**Figure 1.**
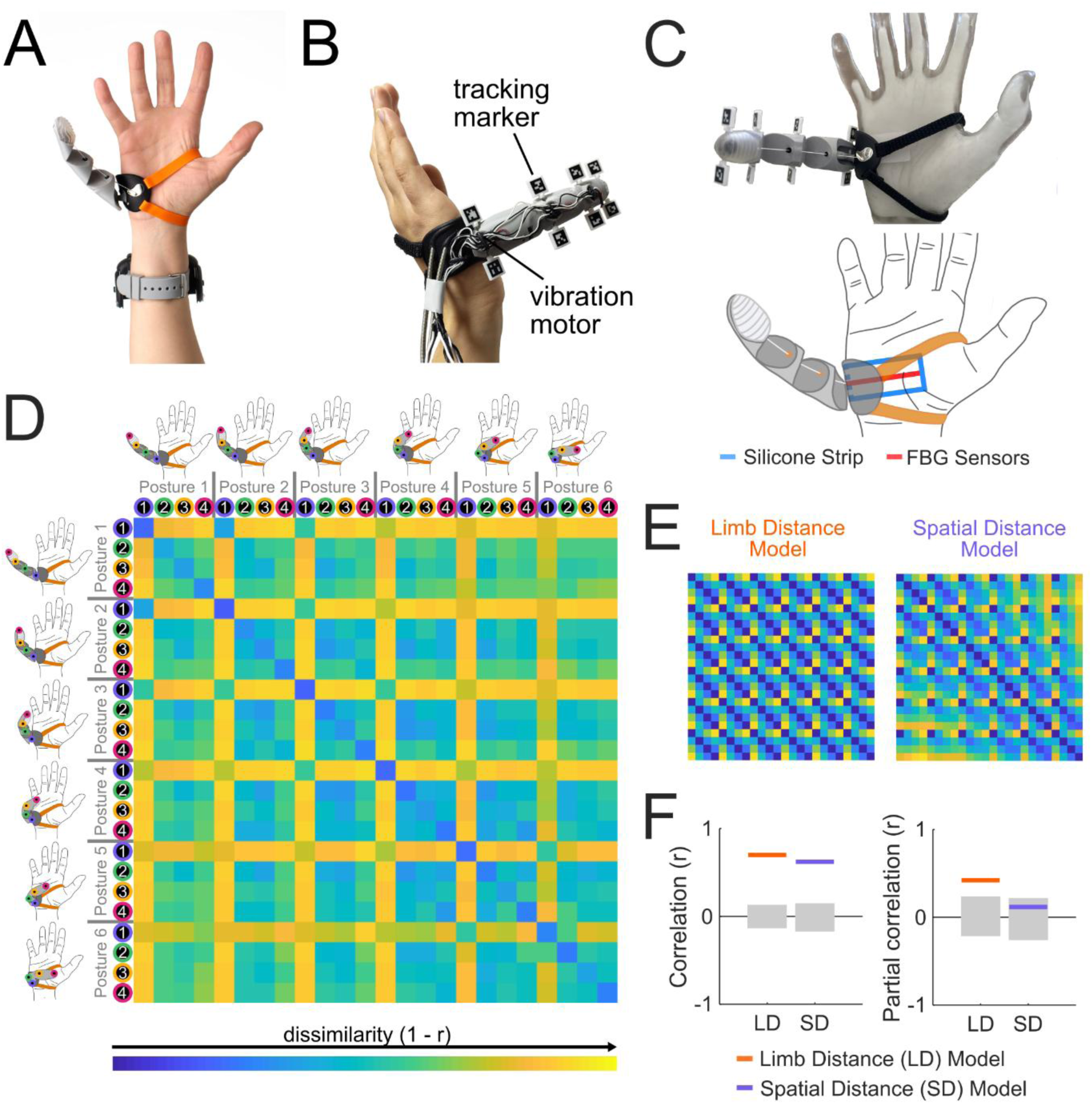
Indirect touch recordings using photonic Fiber Bragg Grating (FBG) sensors. **(A)** The Third Thumb (Dani Clode Design) consists of a soft 3D-printed digit mounted on a rigid plastic handpiece and secured to the hand via two straps positioned above and below the biological thumb. Two servo motors, attached via a wrist strap, control the digit’s posture by pulling on wires connected to the digit, acting against the natural tension of the flexible material to produce postural changes. **(B)** A modified version of the Third Thumb with four vibration motors positioned along the Third Thumb (base, directly after first joint, directly after second joint and tip) to deliver touch stimulation to different Third Thumb locations. Tracking markers (Apriltags) are positioned at the location of each vibration motor to track its spatial position. **(C)** Visualisation of the FBG Sensors embedded in the soft, conformable silicone strip, positioned beneath the Third Thumb handpiece. The silicone strip wraps around the side of the hand, covering the front and back of the Third Thumb handpiece. **(D)** Dissimilarity matrix showing the average 1 - *r* correlation for pairs of recordings (across different blocks) between different touch locations and Third Thumb postures. **(E)** Two models of predicted distances between recordings, one based on the distance of touch locations along the Third Thumb regardless of posture (Limb Distance Model) and one based on the spatial distances of touch locations, taking into account posture (Spatial Distance Model). **(F)** Correlations (left panel) and partial correlations (right panel) of the Limb and Spatial Distance Models with the indirect touch recordings. Grey bars show the 95% confidence interval of the null distribution determined by permutation testing. Both Limb and Spatial Distance Models correlate with the indirect touch recordings. Partial correlations, controlling for the variables in the alternative model, show higher partial correlations for the Limb Distance Model than the Spatial Distance Model.

We first quantified the indirect touch signals reaching the skin to assess whether they contain distinct location-specific information. We then examined whether participants could perceptually discriminate these signals and, importantly, whether they could use them to localise touch in external space by incorporating information about the limb’s posture. Previous work has shown that motor training with robotic limbs leads to improvements in motor control (Amoruso et al., 2022; Dowdall, Molina-Sanchez, et al., 2025; Kieliba et al., 2021; Molina-Sanchez et al., 2026), which may be driven by changes in sensorimotor processing. We therefore tested whether touch localisation abilities develop or improve after motor training with a robotic limb. We found that indirect touch signals provided distinct information about touch location on the Third Thumb and that participants could discriminate these signals and remap them into external space, thus establishing that indirect touch can serve as a functional source of haptic feedback for augmentative artificial limbs.

## 2. Results

### 2.1. Indirect touch signals reflect limb-based touch location distances

To determine whether indirect touch signals convey meaningful information about touch locations, we performed recordings at the Thumb-hand interface using photonic sensors (**Figure S1**). We used a soft, conformable silicone strip equipped with 12 photonic Fiber Bragg Grating (FBG) sensors embedded along an optical fibre and equally spaced within the strip (Massari et al., 2020). The silicone strip was placed horizontally between the Third Thumb handpiece and a custom-designed mannequin hand made with a rigid plastic skeleton and soft plastic surface (**Figure 1C**). We recorded from the 12 FBG sensors while stimulating each of the four vibration motors (0.2 s duration at maximum stimulation) in the six Third Thumb postures. We selected 4 FBG sensors that showed signal changes induced by vibration motor stimulation (including all conditions) and cross-correlated trial time series recordings to align recordings (see Section 5.3. for full details).

We correlated each touch stimulation recording trial time series, extracted from the FBG sensors (**Figure S2**), with each other recording trial time series both within and across Thumb postures (**Figure 1D**) except for trials recorded on the same 32-trial recording block. We then compared the resulting indirect touch recordings matrix with a Limb Distance (LD) Model (dissimilarity reflects touch location distances along the length of the Third Thumb; irrespective of posture), and a Spatial Distance (SD) Model (dissimilarity reflects spatial touch location distances, accounting for posture) (**Figure 1E**). Both models correlated with indirect touch recordings (LD: *r* = 0.70, *p* < .001; SD: *r* = 0.70, *p* < .001), however partial correlations revealed only a higher than chance correlation for the Limb Distance Model (LD: *r* = 0.42, *p* < .001; SD: *r* = 0.12, *p* = .145) (**Figure 1F**). This provides indication that the indirect haptic signals arriving to the skin primarily convey touch information relative to the body, and not space. However, indirect touch signals were also more similar for each touch location across repetitions in the same posture as compared to the same touch location recorded across different postures (**Figure S3**), suggesting some postural modulation of indirect touch signals, despite predominant limb-based information.

### 2.2. Touch localisation on an artificial limb

Next, we assessed whether participants could spatially localise touch on the Third Thumb using indirect touch signals, and whether this ability was enhanced by intensive Third Thumb training (Amoruso et al., 2022; Dowdall, Molina-Sanchez, et al., 2025; Kieliba et al., 2021; Molina-Sanchez et al., 2026). Participants completed two touch localisation sessions with the Third Thumb. The Training Group (n=18) completed five days of motor training between the two sessions, which aimed to enhance their motor skills with the Third Thumb (see **Section 5.5**). These motor tasks involved indirect touch conveyed though natural object manipulation (e.g. moving an object to a target location), rather than vibration motor stimulation (as in the touch localisation sessions). The Control Group (n=20) had a comparable time interval between the two touch localisation sessions, but did not do any Third Thumb training.

In each touch localisation session, participants wore a modified version of the Third Thumb with vibration motors placed at four locations to deliver touch stimulation and markers used to track the spatial position of each touch location via a stereocamera (**Figure 1B**). Participants right hand and the Third Thumb was positioned comfortably in front of them on an arm support and The Third Thumb was automatically positioned in one of six postures (steps of extension – flexion) at the start of each block of trials (**Figure 2A)**. Participants wore shutter goggles to occlude their vision of their hands during each trial (Gomez & Snow, 2023), but allowing them to view their hands and the Third Thumb between trials. Participants also wore headphones to deliver beeps indicating task instructions and white noise to mask vibration sounds. In each experimental trial, a vibration stimulation was delivered to the Third Thumb from one of the four vibration motors. Participants were then instructed to point with their left index finger towards the judged origin of the touch stimulus. We recorded participants pointing movements via markers on their left index finger. We calculated the pointing angle error between participants’ actual pointing vector and the correct pointing vector to the touch stimulus in 3D space, as well as the minimum distance between their pointing vector and the touch stimulus, and the endpoint of their pointing vector on a 2D plane (**Figure 2B**). Pointing endpoints on the 2D plane for one example participant are shown in **Figure 3A**, for two of the six Third Thumb postures.

**Figure 2.**
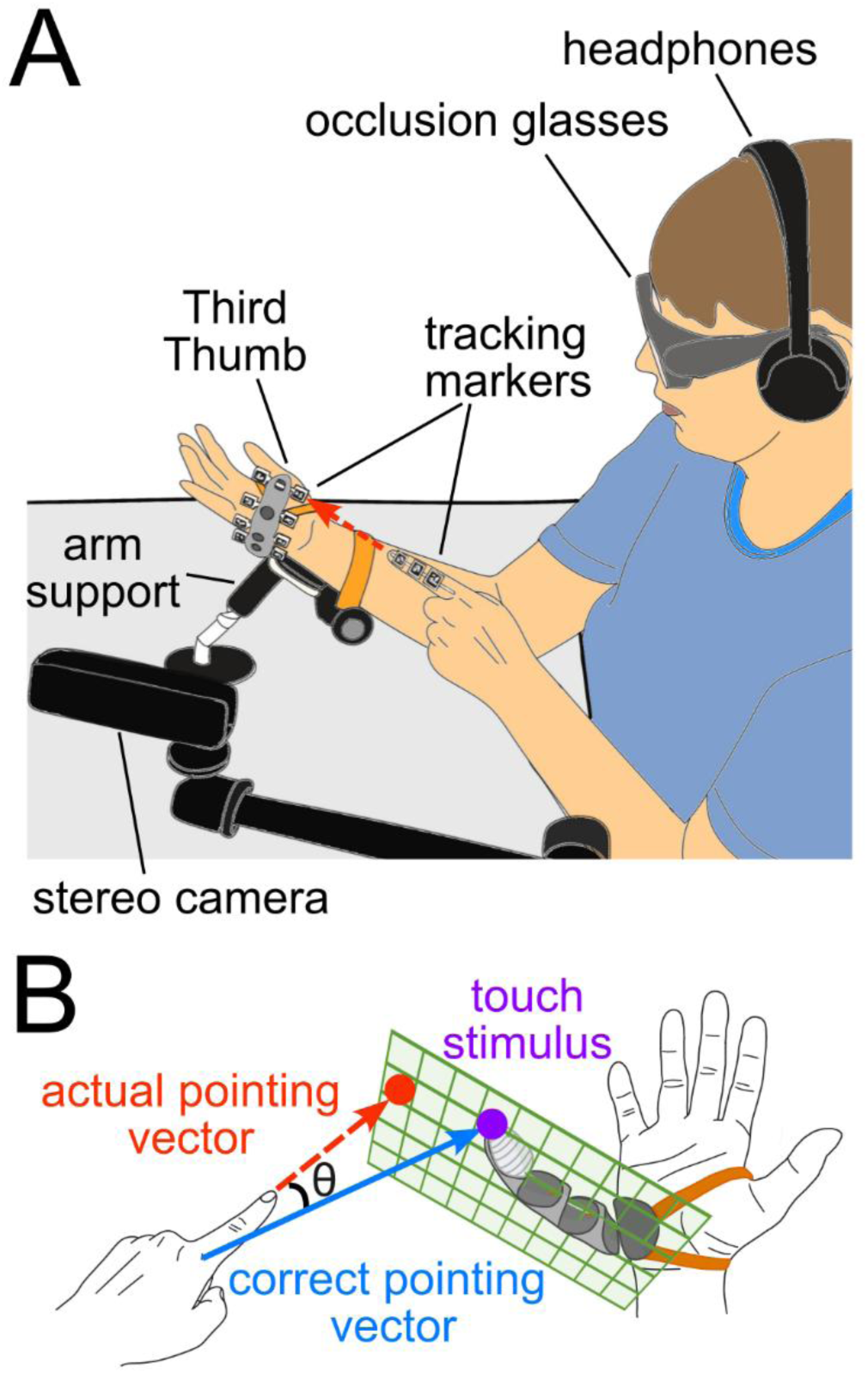
Behavioural Experiment Design. **(A)** Experimental setup. Participants’ wore the Third Thumb on their right hand (Thumb shown in a flexed position towards the palm), which was positioned at a comfortable angle directly in front of them using a custom designed arm support. A stereocamera tracked the 3D positions of tracking markers on the Third Thumb and participants’ left index finger. Participants wore headphones to deliver trial instruction beeps and mask the vibration motor noise and occlusion glasses to hide the position of their hands and the Third Thumb during each trial. The dashed red line indicates the pointing vector following the direction of the participants’ left index finger. **(B)** Experimental task. Participants were instructed to point towards the origin location of the touch stimulus. The angle θ shows the pointing error, defined as the angle between the actual pointing vector (red dashed line) and the correct pointing vector (blue solid line) directly towards the touch stimulus (purple dot). The two-dimensional plane defined by the vibration motor positions along the Third Thumb is shown in green, with the pointing endpoint on the plane illustrated by the red circle.

**Figure 3.**
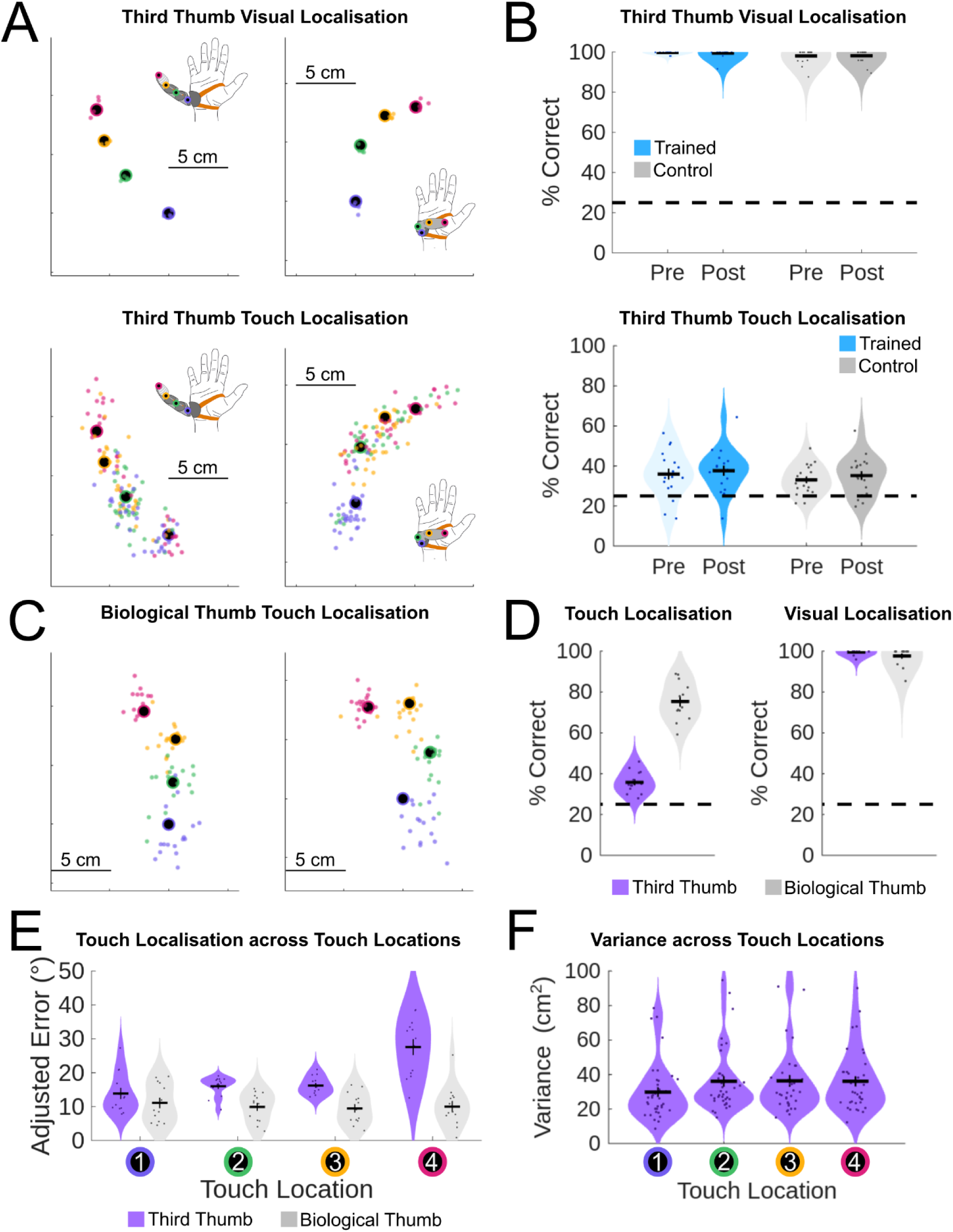
Behavioural touch localisation results for the Third Thumb **(A)** Example participant pointing endpoints on the 2D plane defined by the vibration motor positions along the Third Thumb. Upper panels show visual localisation performance for Third Thumb posture 1 (left) and posture 6 (right), lower panels show touch localisation performance for the same postures. The large circles with black centre show the actual locations of the four touch locations, the individual coloured dots show participants pointing endpoints on individual trials corresponding to each of the four touch locations. **(B)** Percentage correct values for visual (upper) and touch (lower) localisation for the Third Thumb, separated by participant group and session. Correct trials are defined as those where participants’ pointing endpoint was closest to the actual stimulated touch location. The dashed line illustrates chance-level performance and scatter dots show individual participant results. **(C)** Example participant pointing endpoints for touch localisation on the biological right thumb, for two example thumb postures. **(D)** Performance comparison between the Third Thumb and the biological thumb. Percentage correct is shown for touch (left panel) and visual (right panel) localisation. **(E)** Adjusted pointing angle error across touch locations for the Third Thumb and biological thumb. **(F)** Variance in pointing endpoints across touch locations for the Third Thumb.

#### 2.2.1. Spatial localisation of touch on an artificial limb does not require motor training

To validate the experimental setup, participants performed a visual localisation task, where they pointed towards markers positioned at the same four touch locations on the Third Thumb using vision rather than touch cues. Visual localisation endpoints for one example participant are shown in **Figure 3A** (upper panels) for two of six Third Thumb postures. Low performance can indicate technical issues with the setup, or attention drift effects due to task demands. We excluded participants with less than 80% correct accuracy in the visual localisation task from further analyses (n=1). Participants’ accuracy in the visual localisation task (**Figure 3B**, upper panel) was significantly above chance (*V* = 703, *p* < .001, *r* = 1) with correct trials defined as those where participants’ pointing endpoint was closest to the target marker. This demonstrates that our experimental setup can accurately capture participants’ pointing behaviour to locations on the Third Thumb. See **Figure S4A** for further validation using a complementary regression analysis for visual localisation on the Third Thumb. As visual localisation accuracy was close to ceiling performance, we assessed performance differences using non-parametric tests. A Kruskal-Wallis test did not identify any differences in visual localisation performance across experimental groups or pre and post sessions (*H_3_* = 7.1, *p* = .069), indicating that both experimental groups performed similarly across both experimental sessions.

We then assessed participants’ accuracy in the (visually occluded) Third Thumb touch localisation task by calculating whether the closest touch location to their pointing endpoint in each trial was the stimulated touch location or a different touch location. Using this approach, we calculated a percentage correct score for each behavioural session of each participant (**Figure 3B, lower panel**). Both Training Group and Control Group performance was significantly above chance-level (25%) in both pre (Training Group: *t*_17_ = 4.0, *p* < .001, Cohen’s *d* = 1.10; Control Group: *t*_18_ = 4.8, *p* < .001, Cohen’s *d* = 0.95) and post behavioural sessions (Training Group: *t*_17_ = 5.0, *p* < .001, Cohen’s *d* = 1.06; Control Group: *t*_18_ = 4.6, *p* < .001, Cohen’s *d* = 1.17). This result demonstrates that participants were able to successfully perform the touch localisation task with the Third Thumb, even before training. See **Figure S4B** for further validation using a complementary regression analysis for touch localisation on the Third Thumb. Participants’ pointing-based touch localisation accuracy was also comparable to that observed in a perceptual three-alternative forced-choice task (**Figure S5**), suggesting that the additional requirement to integrate limb posture for spatial judgments did not impair performance.

We predicted that participants should improve in performance following Third Thumb training. However, a repeated measures ANOVA showed no main effect of group (*F*(1,35) = 0.8, *p* = .39, *η*_p_^2^ = 0.02), session (*F*(1,35) = 2.6, *p* = .12, *η*_p_^2^ = 0.01) nor interaction between these two factors (*F*(1,35) = 0.0, *p* = .85, *η*_p_^2^ = 0.00). We confirmed this null result using a Bayesian *t*-test of our main contrast of interest (greater improvement in performance across sessions for trained participants compared to controls), which confirmed evidence for the null hypothesis (BF_10_ = 0.28). This suggests that functional experience with the Third Thumb does not contribute to our task demands. As we did not find any changes in performance due to training, our further analyses combine touch localisation data across training groups (n=37) and sessions (n=2) to maximise statistical power.

A subset of participants (n=14) performed an additional experimental session where they performed the same touch localisation task for touch stimuli positioned at comparable locations along their biological right thumb. Pointing endpoints for one example participant in two thumb postures are shown in **Figure 3C**. Participants touch localisation performance for the biological thumb was significantly above chance-level (*t*_12_ = 19.4, *p* < .001, Cohen’s *d* = 5.38), and significantly higher than performance for the Third Thumb (*t*_12_ = 9.03, *p* < .001, Cohen’s *d* = 2.41). **Figure 3D** shows participants’ localisation performance for the biological thumb in comparison to their performance for the Third Thumb.

To assess whether participants’ behavioural performance differed across touch locations and postures for the Third Thumb and the biological thumb we analysed participants’ adjusted pointing angle error (calculated as 3D pointing angle error during touch localisation minus 3D pointing angle error during visual localisation), minimum distance between participants’ pointing vector and the touch location, and their pointing variance. Full results of these analyses are provided in the Supplemental Results. All metrics were lower for the biological thumb as compared to the Third Thumb (**Figure S6A**), indicating that performance was better for the biological thumb than the Third Thumb.

Adjusted pointing angle error and minimum distance were lower for more proximal Third Thumb locations compared to more distal locations (**Figures 3E & S6B & Table S1-2**), and pointing variance was lower for Touch Location 1 compared to all other locations (**Figure 3F & Table S3**). When comparing Touch Location differences for the Third Thumb to the biological thumb, 4 (Touch Location) x 2 (Limb Type) repeated measures ANOVAs identified significant interaction between the two factors for both adjusted pointing angle error and minimum distance. Behavioural performance for Touch Location 1 was comparable for the Third Thumb and biological thumb for both metrics (**Figures 3E & S6B**), suggesting that touch localisation performance for the base of the Third Thumb was comparable to the biological body. Adjusted pointing angle error and minimum distance also varied across Third Thumb postures, with better performance for flexed postures compared to extended ones (**Figure S6C & Table S4-5**).

### 2.3. Representational similarity analysis modelling of participants behaviour: touch localisation reflects spatial coding of touch

We compared participants’ behavioural touch localisation estimates to three different models (**Figure 1**) to determine which best explained localisation responses for the Third Thumb and biological thumb. The Limb Distance Model (LD) predicted that participants’ responses would reflect the distances between touch locations, without linking them to their spatial position. The Spatial Distance Model (SD) predicted that participants would distinguish touch locations along the Third Thumb and integrate this information with the current posture to localise the spatial position. The Touch Input Model (TI) predicted that participants’ responses would reflect the touch input on the skin surface. For the Third Thumb this corresponded to the indirect touch signals reaching the interface between the Third Thumb handpiece and the skin, for the biological thumb the model predicted that touch inputs were distinct for each skin location and independent of one another. **Figure 4** shows the dissimilarity in participants’ behavioural judgements for the different conditions.

**Figure 4.**
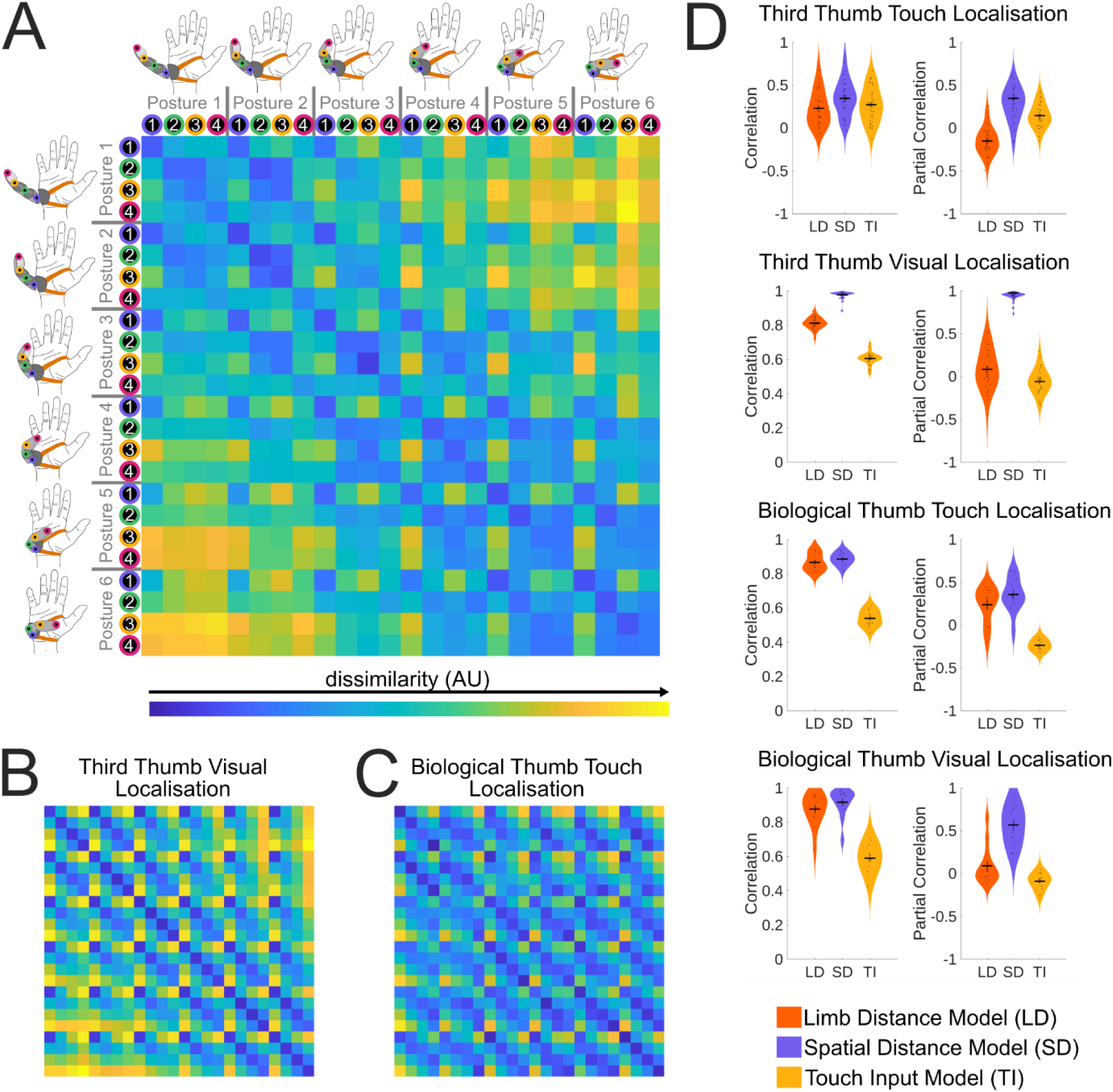
Correlation of touch and visual localisation behavioural performance with Limb Distance, Spatial Distance and Touch Input Models. (A) Third Thumb touch localisation dissimilarity matrix showing the average spatial distances between the mean endpoint on the 2D plane for each condition, averaged across all participants. (B) Third Thumb visual localisation dissimilarity matrix showing the average spatial distances between the mean endpoint on the 2D plane for each condition, averaged across all participants. (C) Biological thumb touch localisation dissimilarity matrix for one example participant. (D) Correlations and partial correlations of touch and visual localisation ability with the Limb Distance Model (LD), the distance of the touch locations along the limb regardless of posture, Spatial Distance Model (SD), the distance between touch locations taking into account posture, and Touch Input Model (TI), the dissimilarity between touch stimulation input on the skin.

Figure 4D shows results of correlations and partial correlations of the LD, SD and TI models with touch and visual localisation judgements for the Third Thumb and biological thumb. All models showed significant correlations with participants’ behavioural judgements, however controlling for variables in the other models using partial correlations revealed differences in model fits to the behavioural data. For touch localisation for the Third Thumb, a repeated measures ANOVA revealed significant differences between model fits (*F*(1.5,55.7)=52.9, *p* < .001, *η*_p_^2^=0.60). This was driven by higher partial correlations for the SD model compared to both other models (SD vs LD: *t*_36_ = 10.1, *p* < .001, Cohen’s *d* = 2.6: SD vs TI: *t*_36_ = 3.2, *p* = .002, Cohen’s *d* = 0.8), and higher partial correlation for the TI model compared to the LD model (*t*_36_ = 1.8, *p* < .001, Cohen’s *d* = 1.8). Partial correlations were positive for the SD and TI models (SD: *t*_36_ = 8.9, *p* < .001, Cohen’s *d* = 1.5; TI: *t*_36_ = 6.3, *p* < .001, Cohen’s *d* = 1.0), but negative for the LD model (*t*_36_ = -5.7, *p* < .001, Cohen’s *d* = 0.9). Thus, participants touch localisation performance for the Third Thumb was best explained by the spatial locations of the touch stimuli, with a contribution of the indirect touch signals reaching the hand.

We compared the Third Thumb touch localisation model fits for to the same three models fit to participants’ visual localisation judgements for the Third Thumb. A repeated measures ANOVA revealed a significant interaction between model partial correlations and localisation condition (*F*(1.9,58.2)=161.8, *p* < .001, *η*_p_^2^=0.20), thus demonstrating that model partial correlations differed between touch and visual localisation. For visual localisation, partial correlations differed across models (*F*(1.6,48.9)=372.6, *p* < .001, *η*_p_^2^=0.93), driven by higher partial correlations for the SD model compared to both other models (SD vs TI: *t*_30_ = 25.2, *p* < .001, Cohen’s *d* = 6.3; SD vs LD: *t*_30_ = 21.6, *p* < .001, Cohen’s *d* = 5.4), and higher partial correlation for the LD model compared to the TI model (*t*_30_ = 3.6, *p* < .001, Cohen’s *d* = 0.9). Partial correlations were positive for the SD model (*t*_30_ = 98.7, *p* < .001, Cohen’s *d* = 17.7), but there was no significant positive or negative partial correlation for the other two models. This result confirms that visual localisation is best modelled by the spatial locations of the stimuli, in contrast to touch localisation with is best modelled by a combination of spatial locations and indirect touch signals.

To determine whether participants touch localisation behaviour for the Third Thumb is similar or distinct to their touch localisation behaviour for the biological thumb, we compared model partial correlations for the same three models with the two different limb types. A 2 x 3 repeated measures ANOVA revealed a significant interaction between limb type and model (*F*(1.2,13.4)=29.7, *p* < .001, *η*_p_^2^=0.53), demonstrating differences in model fits depending on limb type. For the biological thumb, partial correlations differed between models (*F*(1.2,13.0)=33.3, *p* < .001, *η*_p_^2^=0.75), driven by higher partial correlations for the SD model and LD model compared to the TI model (SD vs TI: *t*_11_ = 7.7, *p* < .001, Cohen’s *d* = 3.5; LD vs TI: *t*_11_ = 6.2, *p* < .001, Cohen’s *d* = 2.8). Partial correlations for the SD and LD models did not significantly differ (*t*_11_ = 1.6, *p* = .134, Cohen’s *d* = 0.7). This result shows that the spatial location of the touch stimulation contributes strongly to touch localisation for both the biological thumb and Third Thumb, demonstrating a spatial remapping to limb posture for both limb types. However, different factors also contributed to behavioural performance for the two limb types. For the biological thumb there was a contribution of limb distance, suggesting that prior knowledge of phalange lengths may contribute to participants’ behaviour. In contrast, for the Third Thumb performance was modulated by the incoming indirect touch signals at the wearable interface.

## 3. Discussion

In this study, we investigated whether indirect touch can convey information about the location of tactile stimulation on an artificial limb, the Third Thumb. Our results demonstrate that indirect touch provides distinct signals corresponding to different locations along the limb. Participants were able not only to discriminate these signals but also to map them onto the varying spatial configuration of the Third Thumb arising from postural changes of the soft, flexible Third Thumb digit. This ability indicates a process of multisensory integration, in which tactile cues are combined with information about the limb’s spatial posture provided by visual memory and/or somatosensation. These findings highlight the adaptability and flexibility of the human somatosensory system to process indirect touch signals for artificial body parts.

Our findings demonstrate that the somatosensory system can discriminate indirect touch signals transmitted through an artificial limb with heterogeneous material properties and undergoing posture-dependant geometric transformations. Beyond tactile discrimination, participants’ performance was influenced by the limb’s current spatial configuration, indicating an ability to integrate multisensory cues for an artificial limb. This process likely involves an initial encoding of touch location in limb-centred coordinates, followed by remapping into external space using available multisensory cues indicating limb posture, as has been shown for tools (Themelis et al., 2026). Body posture cues for the Third Thumb may be provided by both somatosensory information and visual memory. Indirect touch has been shown to convey information about artificial limb posture and movement relative to the biological fingers (Kieliba et al., 2021; Molina-Sanchez et al., 2026), thus may provide posture information for remapping. In the present task, participants were able to view the Third Thumb between trials, and thus likely utilised prior visual information to estimate limb posture. Neuroimaging work has demonstrated strong coupling between visual and proprioceptive signals in representing biological body posture (Limanowski & Blankenburg, 2016) which may similarly support encoding of artificial limb posture. Nevertheless, it is important to emphasise that in our design the participant’s hand was visually occluded during the tactile task, and as such the integration of tactile information concerning the tactile target’s location along the Thumb with the visual memory of the Thumb’s position requires a more contextual (and possibly higher-level) process. Altogether, these findings offer new insights for robotics, highlighting the flexibility of the somatosensory system and its potential to support more effective sensory feedback for the motor control of artificial limbs.

Participants exhibited above-chance touch localisation for the Third Thumb even prior to any experience using it, and this performance did not improve following motor training (see also Dowdall, Molina-Sanchez, et al., 2025 for related findings). This suggests that the somatosensory system is already able to process and transform indirect touch signals, potentially through prior experience with tool use. Consistent with this, previous studies have demonstrated accurate touch localisation along tools in the absence of specific training (Miller et al., 2018; Peviani et al., 2026). In mammals, haptic feedback from whiskers is similarly used to infer object properties, pointing to a possible evolutionary basis for this somatosensory capacity (Kleinfeld & Deschênes, 2011). Alternatively, the lack of improvement with training may be because the indirect touch signals in our touch localisation experiment did not closely resemble object interactions during the motor tasks in the Third Thumb training. If so, experience-dependent learning may have been limited to only the specific tactile interactions encountered during training. It is also possible that the lack of changes in touch localisation with training reflects the fact that improvements in Third Thumb motor performance do not directly associate with indirect touch experience. This is supported by the finding that participants show similar learning gains with the untrained hand after training, suggesting that improvement in motor performance can occur independently of haptic experience (Molina-Sanchez et al., 2026). Taken together, these findings suggest that the ability to discriminate, spatially integrate, and transform indirect touch signals is an established feature of the somatosensory system, likely shaped during development and generalised across contexts.

Our indirect touch recordings demonstrate that these signals provide distinct information about touch location along an artificial limb, despite pose-dependent deformations due to posture changes, complex shape, and heterogeneous material properties. Complementing this, our behavioural results confirm that the human somatosensory system can effectively discriminate and utilise these signals. Together, these findings establish indirect touch as a viable and abundant source of tactile feedback for artificial limbs and tools with greater structural complexity than simple rigid, uniform-material devices. These insights have important implications for the design of future artificial limbs. In particular, materials could be selected or engineered to optimise the transmission of distinct indirect tactile signals, thereby enhancing perceptual discriminability. Similarly, the positioning of wearable interfaces could be strategically optimised to maximise effective signal transmission to the user. Altogether, leveraging indirect touch in this way offers a promising route to improving haptic feedback, ultimately supporting more effective sensorimotor integration and closing the sensorimotor control loop in artificial limb systems.

## 5. Methods

### 5.1. Participants

A total of 38 participants (24 women, 15 men, mean age = 26.0 years old ± 5.9, all right handed) took part in the study. From this total sample, the Third Thumb Training Group consisted of 18 participants (13 women, 6 men, mean age = 26.4 years old ± 5.6) who completed two sessions with the Third Thumb with 5 days of Third Thumb training in-between the two sessions (days between sessions: median 7 days, mean 11 days, range 3-56 days). The date of the second session was a median 0 days, mean 1.4 days and range 0-6 days after the last Third Thumb training session. One participant in the Third Thumb Training Group completed their first session after one day of training due to a technical problem.

The Control Group consisted of 20 participants (11 women, 9 men, mean age = 25.6 years old ± 6.3) who completed two sessions with the Third Thumb with a comparable time interval in-between sessions (days between sessions: median 10 days, mean 10.5 days, range 8-15 days). One additional participant dropped out after completing only one experiment session, and their data was not included in the analysis.

14 participants (10 women, 4 men, mean age = 27.5 years old ± 6.6) took part in an additional Biological Thumb session following the Third Thumb sessions. 6 participants were from the Third Thumb Training Group and 8 participants were from the Control Group. The time between the first Third Thumb session and the Biological Thumb Session ranged from 63-255 days with a median of 170 days.

All participants gave informed consent to take part in the study, and the study was approved by the Cambridge Psychology Research Ethics Committee.

### 5.2. The Third Thumb

The Third Thumb (Dani Clode Design) is an additional robotic digit positioned on the side of the hand, with two degrees of freedom controlled by force sensors under the left and right big toes (Figure 1A). The Third Thumb consists of a soft 3D-printed digit that is mounted on a rigid plastic handpiece and secured to the hand via two straps positioned above and below the biological thumb. Two servo motors, attached via a wrist strap, control the digit’s posture by pulling on wires connected to the digit, acting against the natural tension of the flexible material to produce postural changes.

For the touch localisation experimental sessions, only one degree of freedom (flexion – extension) was used and this was controlled automatically rather than by the wearer. A modified version of the Third Thumb was used for the current study (Figure 1B). Four vibration motors were positioned along the Third Thumb (base, directly after first joint, directly after second joint and tip) to deliver touch stimulation to different Third Thumb locations. Tracking markers (Apriltags) were positioned at the location of each vibration motor to track each motors’ spatial position via a stereo camera.

### 5.3. Indirect touch recordings

We recorded the indirect touch signals reaching the interface between the Third Thumb handpiece and the side of the hand when stimulating the same four vibration motors as in the touch localisation experiment sessions. We used a soft, conformable silicone strip (Dragon Skin 10 Medium silicone matrix) containing 12 photonic Fiber Bragg Grating (FBG) sensors embedded along an optical fibre and equally spaced within the strip (Massari et al., 2020) (**Figure S1**). The strip was positioned horizontally between the Third Thumb handpiece and a mannequin hand (Figure 1C). The mannequin hand was 3D printed with a rigid skeleton and softer surface material, simulating the shape and softness of the human hand to provide an optimal fit. Altogether we recorded 3072 indirect touch signal recording trials, consisting of 128 repetitions for each of the four vibration motors and six postures, recorded at a rate of 500 Hz. Each vibration motor was stimulated at maximum for a duration of 0.2 s in each trial.

We processed the indirect touch recording data to identify sensor channels that showed activity changes during vibration motor stimulation and discard inactive sensors outside of the area receiving indirect touch signals. We selected channels that showed a mean absolute change in amplitude of at least 0.0015 nm, in baseline-corrected wavelength shift (Δλ), during vibration motor stimulation (averaged across all vibration motors and postures), resulting in selection of the four central channels. We removed the mean baseline from recordings and aligned indirect touch recordings from different trials using cross-correlation.

We correlated each recording trial with every other trial, expect for trials that were recorded in the same recording block (32 trials per block, 8 repetitions of four vibration motors). We performed correlations separately for each of the four channels and then averaged the correlation across the channels. We then calculated 1 minus the mean correlation for each condition type (four vibration motors, six postures) to calculate a dissimilarity matrix for the indirect touch recordings (Figure 1D). We then compared the indirect touch dissimilarity matrix to two potential models of condition dissimilarity, the Limb Distance Model and Spatial Distance model using correlations and partial correlations. We assessed model fit using permutation testing, where condition labels were permuted 10000 times and the same analysis was performed to create a null distribution of resulting correlations and partial correlations.

### 5.4. Third Thumb touch localisation

Each participant completed two experimental sessions on separate days. The Third Thumb Training Group completed 5 days of Third Thumb training between the two sessions and the Control Group had a similar number of days between sessions.

#### 5.4.1. Experimental setup

The experimental setup is illustrated in Figure 2A. Participants wore the Third Thumb on the side of their right hand. They were seated in front of a table and their right arm was positioned at a comfortable angle using a custom designed arm support. Participants wore headphones to deliver beep sounds indicating the trial beginning and end and white noise to mask vibration motor noise and custom-built occlusion glasses to visually occlude the hands during each trial (Gomez & Snow, 2023). A small rigid rod with three Apriltag markers was taped to the participants’ left index finger and used to record their pointing movements during the experiment. A stereocamera (Luxonis OAK-D Pro) was positioned to the side of the participant and used to detect participants pointing movements and the position of the vibration motors on the Third Thumb using a custom-developed real-time marker-based motion tracking system. The experiment was programmed using Psychtoolbox Version 3.0.19 (Kleiner et al., 2007) with Matlab R2023b using Ubuntu 22.04.4 on a Dell Precision 7680 laptop.

#### 5.4.2. Experiment procedure

Each experiment session consisted of 12 touch localisation blocks, with two blocks for each of the six Third Thumb postures. The order of postures was counterbalanced across participants. Before each pair of blocks, the Third Thumb was moved automatically into the block posture. The stereocamera was adjusted to provide a clear view of all tracking markers. The participant was then instructed to perform a visual localisation procedure, where they pointed towards the four Apriltag markers on the left of the Third Thumb (two repetitions of four markers). This was used to assess tracking and to assess visual localisation ability. Before the first touch localisation block, participants were given a few practice trials to familiarise them with the trial procedure.

Each touch localisation block consisted of 32 trials, with 8 repetitions for each of the four touch stimulation locations, presented in a random order. Participants began each trial with their left index finger touching a metal cylinder positioned to the left of their right arm. A low pitched beep and occlusion of their vision indicated the start of the trial to the participant. Between 0.4 and 1.4 s into the trial, white noise was played via the headphones to mask the vibration motor sounds. One of the four vibration motors was activated at 0.8 s for 0.2 s duration. Participants were instructed to point towards the location they judged to be the origin of the touch stimulation and to hold their finger in position until they heard a high pitched beep indicating the end of the trial at 4.5 s and the glasses becoming transparent, allowing them to view both of their hands and the Third Thumb. Participants then returned their finger to the start position before the next trial began 3.5 s later. A subset of 7 participants (5 Control Group, 2 Training Group) performed the experiment with closing and opening their eyes at the start and end of trials, rather than their vision being automatically occluded by the occlusion glasses.

### 5.5. Third Thumb Training Paradigm

The Third Thumb training involved participants performing a range of tasks on each day that allowed them to improve their motor control of the Third Thumb and its collaboration with their biological fingers to achieve the task goals (Kieliba et al., 2021; Molina-Sanchez et al., 2026). Seven distinct training tasks were implemented: (1) proportional control during transport of an “egg” object requiring precise grip force; (2) expansion of grasp to manipulate multiple objects simultaneously; (3) individuation of the Third Thumb to transport peg-shaped objects; (4) coordinated manipulation of wooden blocks using the Third Thumb and biological fingers to construct a tower; (5) coordinated placement of ring-shaped objects onto pegs; (6) the same ring-placement task performed under an additional cognitive load; and (7) tip-to-tip opposition of the Third Thumb with each biological finger.

All tasks were initially performed in a seated position for one block each. Tasks (3), (4), and (5) were subsequently repeated in a standing position. In total, the training protocol comprised 10 blocks per day, each consisting of five 1-minute trials, resulting in 50 minutes of training per day.

### 5.6. Biological thumb experimental session

14 participants completed an additional experiment session where they performed the same touch localisation experiment for locations on their right biological thumb. The experimental setup differed for this session as follows. Four vibration motors with Apriltag markers above them were taped to four locations along the biological thumb: the carometacarpal (CMC) joint, just above the metacarpophalangeal (MCP) joint, just above the interphalangeal (IP) joint and at the thumb tip (palm side, away from the nail). These locations correspond to the positions of the vibration motors on the Third Thumb. A custom designed thumb rest was used to support the participants arm and vary the angles of the thumb joints across postures. Six different combinations of two support angles were used to vary thumb posture, however the resulting thumb position also depended on participants’ thumb size and comfort. The experiment procedure was the same for the biological thumb session as for the Third Thumb session.

### 5.7. Behavioural data processing and statistical analyses

We recorded participants’ pointing angle error and pointing endpoints on a 2D plane defined by the vibration motor positions. Pointing angle error was calculated as the angle between the participants’ actual pointing vector and the vector from the lowest marker on their left index finger directly towards the stimulated touch location (Figure 2B). For each touch localisation trial we selected the median pointing angle and endpoint in a 2 – 3.4 s time window following the touch stimulus. This time window ensured participants’ had stable pointing towards the judged touch stimulus origin location and ended shortly before the 3.5 s beep signalling the trial’s completion to minimise any effects of early anticipation of the trial end. For visual localisation, we used a 3 – 5.5 s time window following the marker instruction, to ensure participants’ had stable pointing and to minimise any effects of early anticipation of the trial end at 6 s. We removed outliers prior to data analysis by removing trials where the pointing endpoint was greater than 15 cm from any touch location and trials where the pointing endpoint or pointing angle error was more than three standard deviations from the mean for each condition.

We calculated percentage correct values for each participant by defining correct trials as trials where the closest touch location to the participants’ pointing endpoint was the stimulated touch location. We calculated adjusted pointing angle error for each condition as pointing angle error during touch location minus pointing angle error during visual location for each condition, thus accounting for each participants’ pointing accuracy under optimal conditions. We performed statistical analyses using JASP 0.17.1.0. We applied non-parametric tests for data that was not normally distributed and corrected for non-sphericity using Greenhouse-Geisser correction. In cases where multiple tests or post-hoc tests were conducted we corrected for multiple comparisons using Holm-Bonferroni correction.

### 5.8. Representational similarity analysis

We compared participants’ behavioural responses to different models to determine which models could best explain participants’ behaviour in the touch and visual localisation experiments. We calculated dissimilarity matrices for participants’ touch and visual localisation responses for both the Third Thumb and biological thumb, by calculating the distance between mean pointing endpoints for each pair of conditions (4 touch locations x 6 postures). We then compared behavioural response dissimilarity matrices to Limb Distance, Spatial Distance and Touch Input Models of condition dissimilarity. For each dissimilarity matrix, we extracted the unique pairwise comparisons and performed z-score normalisation before performing correlations and partial correlations between behavioural response dissimilarity matrices and models.

## Acknowledgements

This work was supported by a Walter Benjamin Fellowship of the German Research Foundation (DFG) awarded to Celia Foster (project number: 545902796), UKRI’s Frontier Research Guarantee (EP/X040372/1), the Medical Research Council (MC_uu_00030/10) and a Wellcome Trust Senior Research Fellowship (215575/Z/19/Z).

We thank Jasmine Chan, Zehra Merchant, Viktorija Pavalkyte and Ariel Seng for assistance with data collection and participant recruitment, Lucy Dowdall, David Hayes and Mark Townsend for technical advice and assistance, Ema Jugović, Julien Russ, Eden Lechelle, Angus Gray, Ellie Carre and Mabel Ziman for Third Thumb training of participants and Sarah Buehler and Mathew Kollamkulam for pilot touch discrimination studies.

