## Supplemental Information for "Spatial localisation of touch on a robotic limb in the absence of direct haptic feedback"

### Supplemental Material

### Supplemental Results

##### Third Thumb Regression analysis

We performed a regression analysis to determine whether participants' pointing endpoints in the visual and touch localisation tasks was comparable to the actual locations of the stimuli on the Third Thumb. To do this we used the Matlab mvregress function to determine the slopes fitting the x and y coordinates of participants' pointing endpoints. A slope of 1 indicates a perfect fit of the behavioural judgements to the actual spatial touch location positions. For each condition, we additionally calculated a null distribution by permuting condition labels and re-running the regression, with 10000 permutations in total.

For visual localisation (**Figure S4A**), both x-axis and y-axis regression slopes were close to one and significantly higher than the null distribution for both participant groups and experimental sessions ($p$ < .001 for all 4 conditions for both x-axis and y-axis). Results were comparable for touch localisation (**Figure S4B**), with both x-axis and y-axis regression slopes significantly higher than the null distribution for both participant groups and experimental sessions ($p$ < .001 for all 4 conditions for both x-axis and y-axis). Thus, our regression analysis demonstrates that participants were able to successfully perform the touch location task for the Third Thumb.

##### Touch localisation performance differs across stimulation locations and limb posture

We tested whether touch localisation performance differs across touch locations along the Third Thumb or different postures. We assessed differences in adjusted pointing angle error, calculated as the 3D angle of error in participants' pointing towards the touch location target **(Figure 2B)** minus their error when pointing to markers at the same positions along the Third Thumb with vision. Thus, this value assesses performance in our conditions of interest while controlling for individual differences in pointing ability under optimal conditions. We also analysed the minimum distance of participants' pointing vector to the stimulated touch location and the variance of participants pointing endpoints on the 2D plane.

We used a repeated measures ANOVA to test whether adjusted pointing angle errors differ across touch locations and postures, and identified significant main effects for touch location ($F$(2.0,70.9)=27.7, $p$ < .001, $\eta_p^2$=0.44) and posture ($F$(2.7,95.6)=6.9, $p$ < .001, $\eta_p^2$=0.16), as well as significant interaction between these two factors ($F$(5.3,189.6)=10.5, $p$ < .001, $\eta_p^2$=0.23). Follow-up tests showed that performance was worse for more distal touch locations compared to more proximal locations (**Figure 3E and Table S1**) and worse for more extended Third Thumb postures compared to more flexed postures (**Figure S6C and Table S4**).

When considering minimum distance of the pointing vector to the stimulated touch location, we found consistent results. We repeated measures ANOVA identified significant main effects for touch location ($F$(1.8,63.7)=263.5, $p$ < .001, $\eta_p^2$=0.50) and posture ($F$(2.7,96.4)=3.2, $p$ = .031, $\eta_p^2$=0.08), as well as significant interaction between these two factors ($F$(4.5, 161.0)=5.3, $p$ < .001, $\eta_p^2$=0.13). Follow-up tests showed that performance using this metric was worse for Touch Location 4 compared to more proximal locations (**Figure S6B and Table S2**) and worse for Posture 2 compared to Postures 5 and 6 (**Figure S6C and Table S5**).

A repeated measures ANOVA testing whether variance differs across touch locations and postures identified a significant main effect of touch location ($F(2.6,94.9)=13.8$, $p < .001$, $\eta_p^2=0.28$), but no differences across postures ($F(3.4,122.6)=0.4$, $p = .774$, $\eta_p^2=0.01$) or interaction between touch location and posture ($F(6.3,225.5)=1.1$, $p = .379$, $\eta_p^2=0.03$). Follow-up tests showed that variance was lower for touch location 1, the Third Thumb base, compared to all other touch locations (**Figure 3F and Table S3).**

Lastly, we compared performance during touch localisation for the Third Thumb to touch localisation for the biological right thumb. We used a repeated measures ANOVA to test whether adjusted pointing angle error showed differences across Limb Type (Third Thumb, Biological Thumb) and Touch Location. We identified significant main effects for limb type ($F(1,12)=63.1$, $p < .001$, $\eta_p^2=0.37$) and touch location ($F(1.5,18.0)=12.5$, $p < .001$, $\eta_p^2=0.14$), as well as a significant interaction between these two factors ($F(2.0,23.6)=14.0$, $p < .001$, $\eta_p^2=0.16$). Follow-up paired *t*-tests showed that participants performed better for their biological thumb than for the Third Thumb (**Figure S6A**, $t_{12}= 7.9$, $p < .001$, Cohen's d=1.6). Participants did not show differences in behavioural performance across touch locations for the biological thumb (**Figure 3E**, $F(2.1,25.2)=0.8$, $p = .470$, $\eta_p^2=0.06$), in contrast to the differences identified above for the Third Thumb. No differences were found between touch localisation performance for touch location 1 for the Third Thumb and biological thumb, suggesting that touch localisation performance for the base of the Third Thumb was comparable to the biological body ($t_{12}= 1.0$, $p = .353$, Cohen's d=0.27).

Our results were highly consistent for minimum distance. A repeated measures ANOVA identified significant main effects for limb type ($F(1,12)=45.3$, $p < .001$, $\eta_p^2=0.79$) and touch location ($F(1.6,19.3)=20.2$, $p < .001$, $\eta_p^2=0.63$), as well as a significant interaction between these two factors ($F(1.9,22.5)=18.0$, $p < .001$, $\eta_p^2=0.60$). Follow-up paired *t*-tests showed that participants has lower minimum distance for their biological thumb than for the Third Thumb (**Figure S6A**, $t_{12}= 6.7$, $p < .001$, Cohen's d=1.9). Participants did not show differences in minimum distance across touch locations for the biological thumb (**Figure S6B**, $F(1.9,22.4)=0.0$, $p = .967$, $\eta_p^2=0.00$), and no differences were found between touch localisation performance for touch location 1 for the Third Thumb and biological thumb, providing further evidence that touch localisation performance for the base of the Third Thumb was comparable to the biological body ($t_{12}= 1.9$, $p = .084$, Cohen's d=0.52).

For variance, a repeated ANOVA with the same factors identified a significant main effects of Limb Type ($F(1,12)=20.7$, $p < .001$, $\eta_p^2=0.54$) and Touch Location ($F(2.3,28.1)=7.9$, $p = .001$, $\eta_p^2=0.03$), but no interaction between these two factors ($F(2.1,24.8)=2.3$, $p = .124$, $\eta_p^2=0.012$). Follow-up tests showed variance was lower for the biological thumb than the Third Thumb (**Figure S5A**, $t_{12}= 15.1$, $p < .001$, Cohen's d=1.7), and lower for touch location 1 than all other touch locations.

### Supplemental Figures

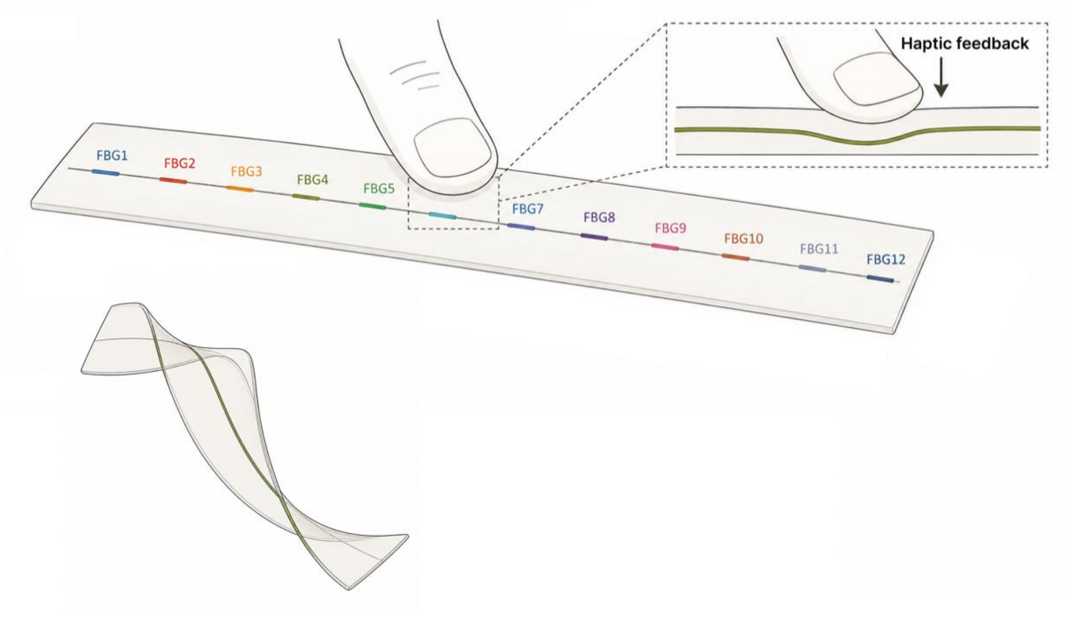


**Figure S1.** Silicone optic fiber strip equipped with 12 photonic Fiber Bragg Grating (FBG) sensors.

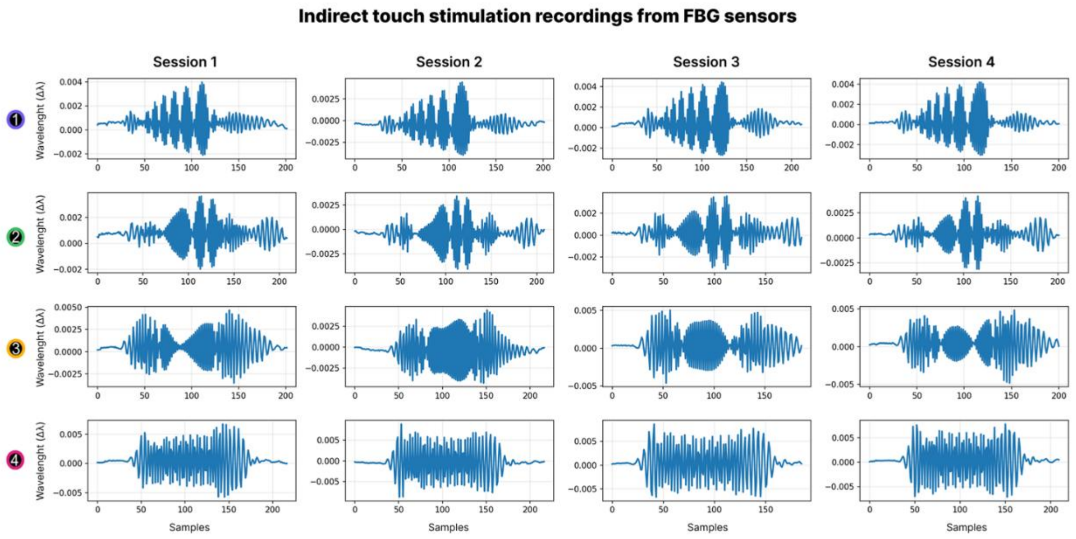


**Figure S2.** Indirect touch stimulation recordings from FBG sensors. Signal recorded from one FBG sensor (number 7) across different sessions (horizontally), and across touch location 1 to 4 (vertically).

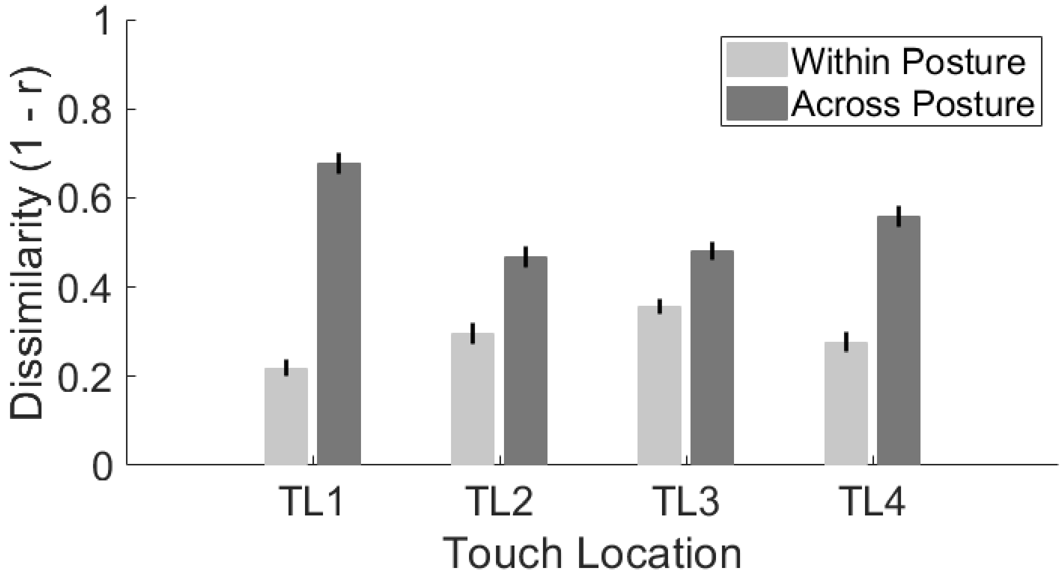


**Figure S3.** Dissimilarity of indirect touch stimulation recordings for the same touch locations within the same posture (light grey bars) and across postures (dark grey bars). Error bars show standard error of the mean. Recordings were more dissimilar across postures as compared to different recordings within the same posture, assessed by a 4 x 2 ANOVA that identified a significant main effect of within vs across posture ($F(1,160) = 188.1$, $p < .001$, $\eta_p^2 = 0.54$).

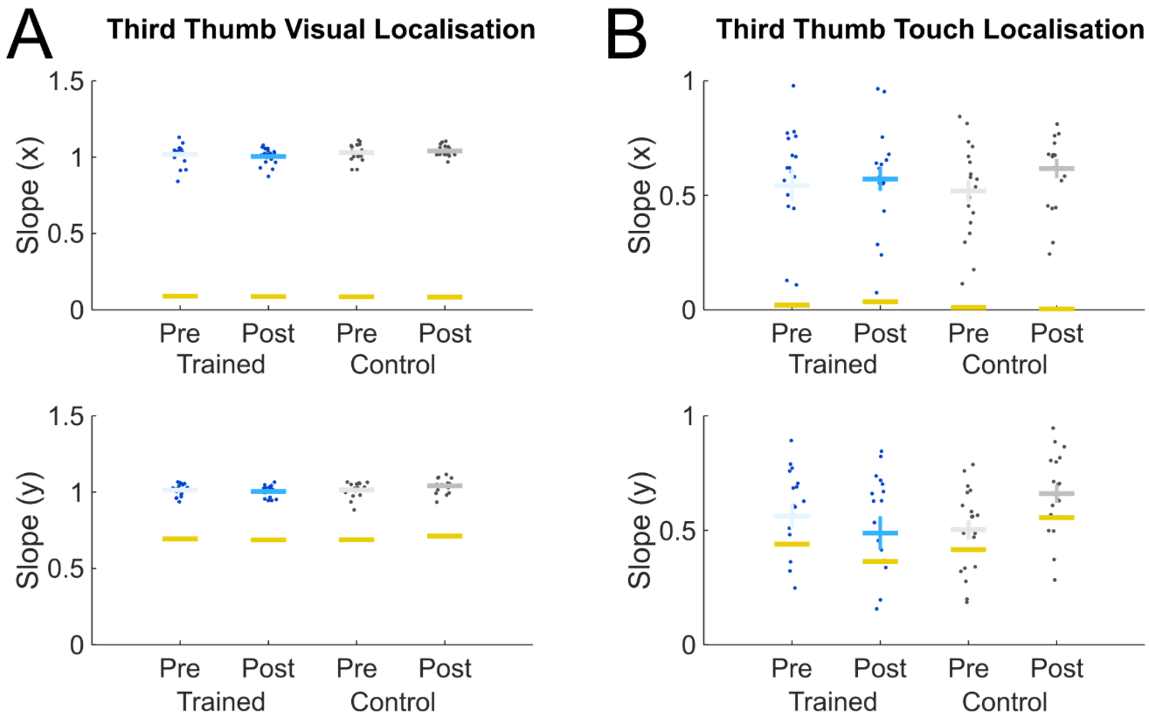


**Figure S4.** Regression analysis for visual **(A)** and touch **(B)** localisation on the Third Thumb. This analysis tested whether participants' pointing endpoints in the visual and touch localisation tasks is comparable to the actual locations of the stimuli on the Third Thumb, by fitting slopes to fit the x and y coordinates of participants' pointing endpoints to the spatial locations of the touch stimuli, with a slope of 1 indicating a perfect fit of the behavioural judgements to the actual spatial touch location positions. Upper plots show regression slopes for the x-axis and lower plots show regression plots for the y-axis slopes across training groups and experiment sessions. Horizontal blue lines indicate mean slopes for the Trained group and horizontal grey lines indicate mean slopes for the Control group. Scatter points indicate individual participant results and error bars show standard error of the mean. Yellow lines indicate the 95$^{th}$ percentile of the null distribution, calculated by permutation testing of participants' data (10000 permutations) and re-running the regression.

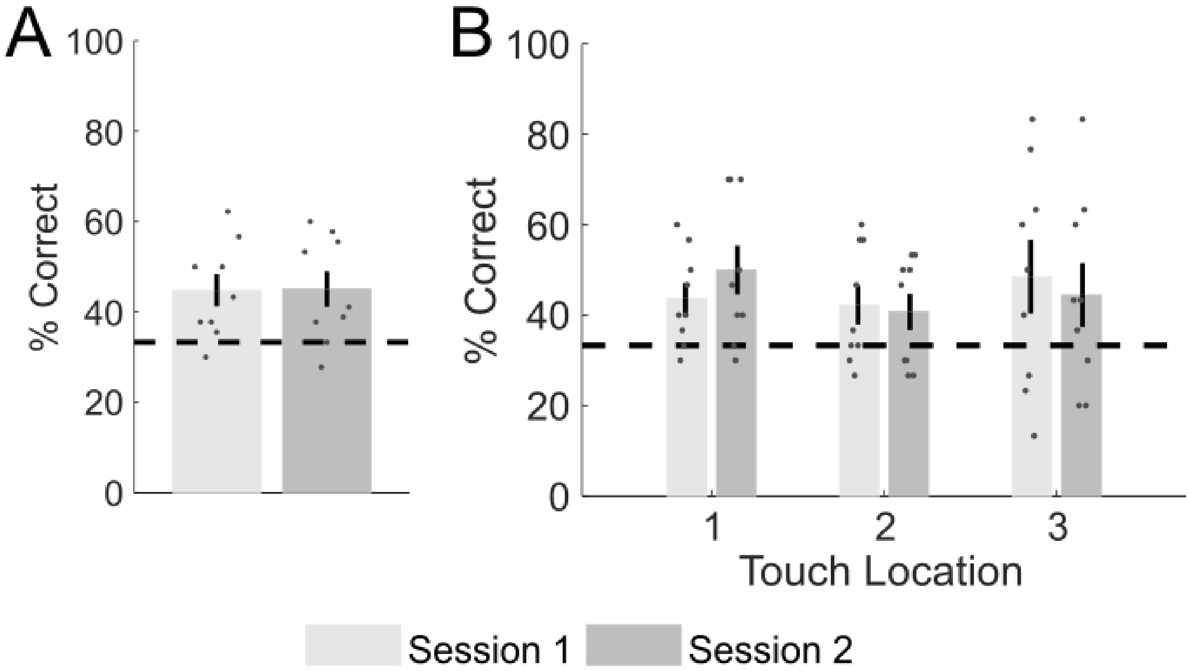


**Figure S5.** Touch discrimination for three touch locations along the Third Thumb. Participants received touch stimulation via vibration motors placed at three locations along the Third Thumb and indicated via a keyboard response which of the three locations they judged was stimulated in each trial (3 alternative force choice). **(A)** Overall touch discrimination performance for n=9 participants. Each participant took part in two experimental sessions on different days. **(B)** Touch discrimination performance for each of the three touch locations (proximal to distal). Bars show group mean performance, error bars show standard error of the mean, dashed lines indicate chance-level performance (1/3) and scatter points show individual participant performance.

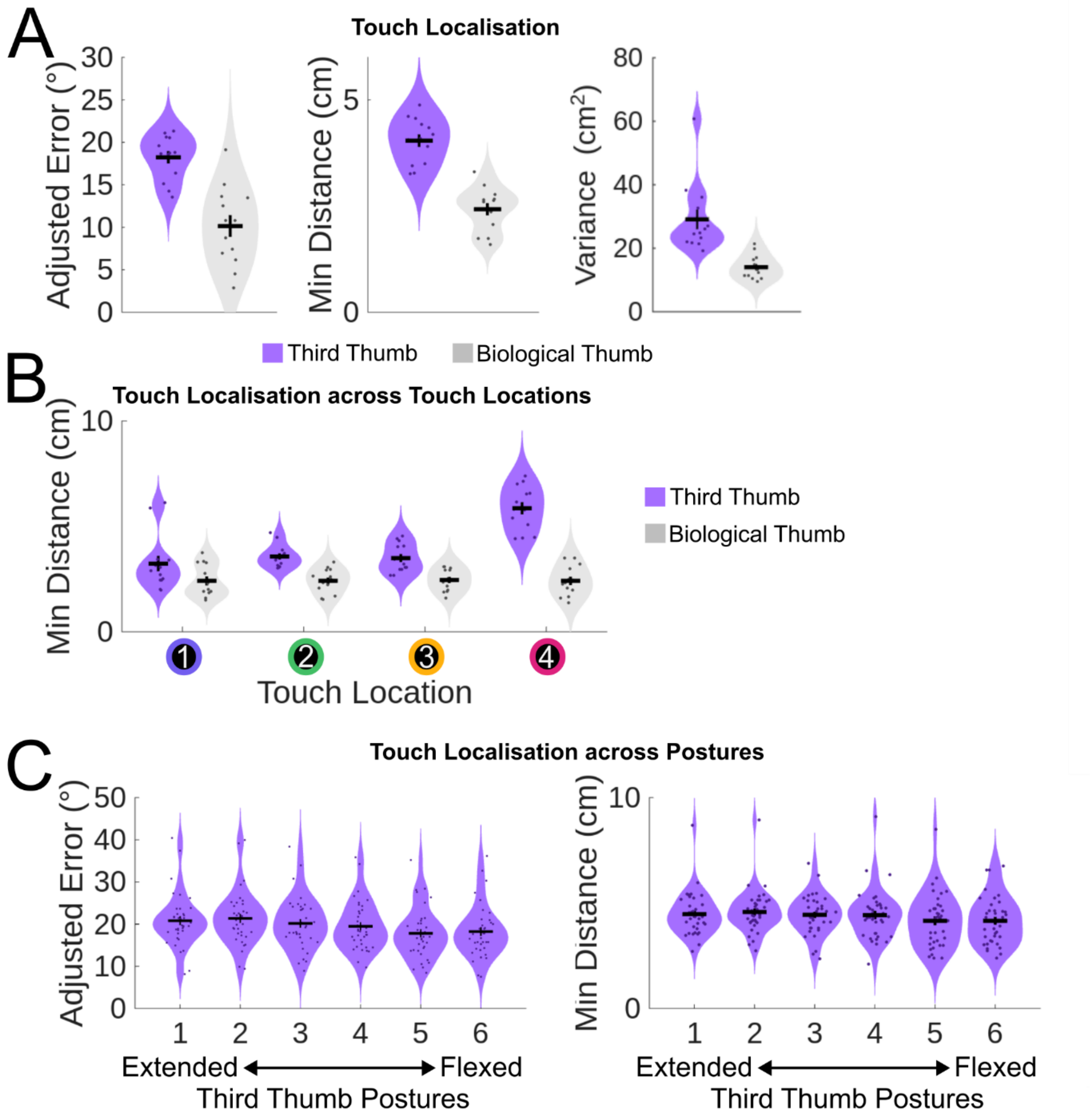


**Figure S6.** Behavioural performance across limb type and Third Thumb postures and touch locations. **(A)** Adjusted pointing angle error, minimum distance of the pointing vector to the touch location, and variance during touch localisation for the Third Thumb and biological thumb. Adjusted pointing angle error is calculated as the pointing angle error during touch localisation minus the pointing angle error during visual localisation. **(B)** Minimum distance across touch locations for the Third Thumb and biological thumb. **(C)** Adjusted pointing angle error and minimum distance across postures for the Third Thumb.

#### Supplemental Tables

| Compared Touch Locations (TL) | Mean Difference (°) | *t* value | *p* value (Holm-Bonferroni corrected) |
|---|---|---|---|
| TL 1 & TL 2 | 6.9 | 4.7 | < .001* |
| TL 1 & TL 3 | 9.9 | 6.7 | < .001* |
| TL 1 & TL 4 | 12.9 | 8.7 | < .001* |
| TL 2 & TL 3 | 3.0 | 2.0 | .091 |
| TL 2 & TL 4 | 5.9 | 4.0 | < .001* |
| TL 3 & TL 4 | 3.0 | 2.0 | .091 |

**Table S1.** Comparison of adjusted pointing angle error across different touch locations on the Third Thumb. * indicates significant results.

| Compared Touch Locations (TL) | Mean Difference (cm) | *t* value | *p* value (Holm-Bonferroni corrected) |
|---|---|---|---|
| TL 1 & TL 2 | 0.1 | 0.3 | .783 |
| TL 1 & TL 3 | 0.6 | 2.2 | .097 |
| TL 1 & TL 4 | 2.3 | 9.1 | < .001* |
| TL 2 & TL 3 | 0.5 | 1.9 | .123 |
| TL 2 & TL 4 | 2.3 | 8.8 | < .001* |
| TL 3 & TL 4 | 1.7 | 6.9 | < .001* |

**Table S2.** Comparison of minimum distance of the pointing vector to the stimulated touch location across touch locations on the Third Thumb. * indicates significant results.

| Compared Touch Locations (TL) | Mean Difference ($cm^2$) | *t* value | *p* value (Holm-Bonferroni corrected) |
|---|---|---|---|
| TL 1 & TL 2 | -6.2 | -5.2 | < .001* |
| TL 1 & TL 3 | -6.5 | -5.4 | < .001* |
| TL 1 & TL 4 | -6.2 | -5.2 | < .001* |
| TL 2 & TL 3 | -0.2 | -0.2 | 1.000 |
| TL 2 & TL 4 | 0.0 | 0.0 | 1.000 |
| TL 3 & TL 4 | 0.2 | 0.2 | 1.000 |

**Table S3.** Comparison of pointing variance across touch locations on the Third Thumb. * indicates significant results.

| Compared Postures | Mean Difference (°) | *t* value | *p* value (Holm-Bonferroni corrected) |
|---|---|---|---|
| Pos 1 & Pos 2 | -0.6 | -0.7 | 1.000 |
| Pos 1 & Pos 3 | 0.6 | 0.8 | 1.000 |
| Pos 1 & Pos 4 | 1.3 | 1.7 | 0.606 |
| Pos 1 & Pos 5 | 3.0 | 3.9 | 0.002* |
| Pos 1 & Pos 6 | 2.6 | 3.4 | 0.009* |
| Pos 2 & Pos 3 | 1.2 | 1.6 | 0.610 |
| Pos 2 & Pos 4 | 1.9 | 2.4 | 0.139 |

| | | | |
|---|---|---|---|
| Pos 2 & Pos 5 | 3.6 | 4.6 | < 0.001* |
| Pos 2 & Pos 6 | 3.2 | 4.1 | < 0.001* |
| Pos 3 & Pos 4 | 0.7 | 0.9 | 1.000 |
| Pos 3 & Pos 5 | 2.4 | 3.1 | 0.026* |
| Pos 3 & Pos 6 | 2.0 | 2.6 | 0.106 |
| Pos 4 & Pos 5 | 1.7 | 2.2 | 0.235 |
| Pos 4 & Pos 6 | 1.3 | 1.7 | 0.606 |
| Pos 5 & Pos 6 | -0.4 | -0.5 | 1.000 |

**Table S4.** Comparison of adjusted pointing angle error across Third Thumb postures. * indicates significant results.

| **Compared Postures** | **Mean Difference (cm)** | ***t* value** | ***p* value (Holm-Bonferroni corrected)** |
|---|---|---|---|
| Pos 1 & Pos 2 | -0.1 | -0.8 | 1.000 |
| Pos 1 & Pos 3 | 0.0 | 0.2 | 1.000 |
| Pos 1 & Pos 4 | 0.0 | 0.3 | 1.000 |
| Pos 1 & Pos 5 | 0.3 | 2.3 | .293 |
| Pos 1 & Pos 6 | 0.3 | 2.3 | .293 |
| Pos 2 & Pos 3 | 0.1 | 1.0 | 1.000 |
| Pos 2 & Pos 4 | 0.1 | 1.1 | 1.000 |
| Pos 2 & Pos 5 | 0.4 | 3.0 | .039* |
| Pos 2 & Pos 6 | 0.4 | 3.0 | .038* |
| Pos 3 & Pos 4 | 0.0 | 0.1 | 1.000 |
| Pos 3 & Pos 5 | 0.3 | 2.0 | .439 |
| Pos 3 & Pos 6 | 0.3 | 2.1 | .439 |
| Pos 4 & Pos 5 | 0.3 | 2.0 | .439 |
| Pos 4 & Pos 6 | 0.3 | 2.0 | .439 |
| Pos 5 & Pos 6 | 0.0 | 0.0 | 1.000 |

**Table S5.** Comparison of minimum distance of the pointing vector to the stimulated touch location across Third Thumb postures. * indicates significant results.
